# Rhizobial adhesives enhance nodule formation in sunn hemp

**DOI:** 10.1101/2021.09.27.461990

**Authors:** Qulina Rai, Robin Choudhury, Pushpa Soti, Alexis Racelis

**Author notes:** Correspondence: Alex Racelis, The University of Texas Rio, Grande Valley, School of Earth, Environment and Marine Sciences.

## Abstract

Inoculation of legume seed with rhizobacteria before planting is an efficient and convenient way of introducing effective rhizobacteria to soil vicinity of legume root and ensuring nitrogen fixation in cover cropped field. However, there are still challenges in identifying the proper seed coating technique to ensure microbial survival in adverse environmental conditions and maintaining the symbiotic relation with plants. The objectives of this study are firstly, to analyze the effectiveness of different sticking agents aiding inoculation of *Bradyrhizobium japonicum* L. in sunn hemp seeds to enhance root nodule formation. Secondly, to observe nodulation pattern over time as affected by the treatment and lastly to check if there is significant difference between main root and lateral root nodulation pattern due to the treatments. Two similar field studies were conducted in fall 2019 and summer 2020 using four sticking agents: water, peanut oil, 10% jaggery solution, and 40% gum arabic solution. The fall study showed no significant differences among total nodules across treatments, but percentage of active nodules was highest in the oil treatment and lowest in the water treatment. In the summer study, significantly higher total nodules were seen in the jaggery treatment and the lowest was in water treatment again, however, there were no differences in percentage of total active nodules across treatments. Interestingly, the trend across weeks showed gum arabic treatment exhibiting higher main root nodulation and jaggery treatment exhibiting higher lateral root nodulation. Overall, water as an adhesive was less effective in aiding nodulation compared to other treatments. Peanut oil and jaggery showed better performance as adhesives aiding active nodulation and could be more effective than gum arabic or water.

## 1. INTRODUCTION

Cover cropping is a conservation strategy where crops are grown primarily to improve soil health. It improves soil organic matter, soil fertility, water infiltration and retention, as well as suppresses weed emergence and disrupts pest life cycles (Snapp et al., 2005; Soti and Racelis, 2020). Legumes used in cover cropping contribute largely to sustainable nitrogen management through biological nitrogen fixation (Sarrantonio and Gallandt, 2003) and estimated to contribute 50-70 Tg of nitrogen each year to the global nitrogen budget, 18-26 Tg of which is within agricultural systems (Herridge et al., 2008). Presently, many farmers utilize leguminous cover crops such as vetch, cowpea, and sunn hemp for the potential nitrogen gains (Sarrantonio and Gallandt, 2003).

The benefit of biological nitrogen fixation can be harnessed only when appropriate live rhizobacteria in soil contact legume root to form nodules. Rhizobia are Gram-negative soil bacteria with the unique ability to establish a nitrogen-fixing symbiosis on legume roots and on the stems of some aquatic legumes. Though legume-rhizobium partnership is found together in their native habitats, agricultural systems have majority of its legumes depending on inoculant for their desired symbiont (O’Hara, 2001). Whether rhizobia are native or introduced through an inoculant, the initiation of nitrogen fixation starts with bacteria’s infection of the legume root. Rhizobia recognize signal compounds from the roots of host plants and adhere to root hairs beginning to synthesize and export a nodulation factor to stimulate nodule formation process in their hosts (Gage, 2004). Plants then begin curling their root hairs around the rhizobia leading to formation of infection pocket (Murray, 2011). As the rhizobia grow and reproduce, the infection pocket develops into a branching infection thread, which further invade the nodule cells (Gage, 2004; Murray, 2011). On successful invasion and interaction of rhizobia with root hairs, nodule gets further developed and is finally capable of fixing nitrogen.

Occasionally, this interaction between rhizobia and plant root parts fails to form nodules due to several environmental conditions such as soil temperature, soil moisture, salinity, nutrient deficiency, soil pH (Kasper et al., 2019) and competition with native rhizobia (Miller and May, 1991). Additionally, inoculating with the right strain in high numbers that can outcompete the native-naturalized rhizobia under field conditions thus preventing their impact on the inoculated strain is important (Brockwell et al., 1995; Brockwell et al., 1987; Irisarri et al., 2019; Nambiar et al., 1987; Smith et al., 1981; Somasegaran et al., 1988). While there are several inoculation techniques such as dusting, slurry, pelleting, and vacuum impregnation (Deaker et al., 2004), no inoculation technique assures certainty andconsistency in field conditions. Thus, choosing a proper inoculation technique is necessary to ensure the seeds receive the intended rhizobia.

The choice of proper inoculation technique is generally dependent on the environmental characteristics of the site and the specific plant-inoculant species interaction. Effective nodulation of legumes is achieved when the threshold of rhizobial inoculant per seed is achieved, which is 1 × 10^5^ rhizobia/seed for large-seeded, 1 × 10^4^/seed for medium-seeded, and 1 × 10^3^/seed for small-seeded legumes (O’Hara, 2001). Type of adhesive used, and inoculation method affect the attainment of this threshold. Inconsistent inoculant coverage due to poor adhesion often leads to inoculant loss during seeding process. For example, water, one of the most widely used sticking agents, provides good adhesion initially but the inoculant tends to crumble off during handling and seeding process (Hoben et al., 1991). Other sticking agents such as oil, sugar, jaggery, a by-product of sugarcane industry, and gum arabic are reported to have good adhesion with seeds promoting root hair infection by rhizobia (Hoben et al., 1991; Sayyed et al., 2012). Sugar is known to provide good moisture retention which is important for the survival of rhizobia (Hoben et al., 1991). Gum arabic is reported to be effective in a wide range of temperature but is expensive compared to other agents (Hoben et al., 1991). Overall, using vegetable oil as a sticking agent can be reasonably inexpensive, readily available, non-toxic method to inoculate seeds.

Sunn hemp (*Crotalaria juncea*), a legume, has become increasingly popular as a summer cover crop in the topics and subtropics. It produces 5604 kg of biomass/hectare and 135kg/ha N in a 9–12-week period and grows rapidly (Sheahan, 2012) reaching maturation between 60 and 90 days. Sunn hemp is drought tolerant and has potential to build organic matter levels and sequester soil carbon (Sheahan, 2012). It is known to form symbiotic associations with *Bradyrhizobium japonicum* and fixes nitrogen in the root nodules (Kasper et al., 2019). In addition, sunn hemp is reported to be resistant to root knot nematode (McSorley, 1999) and can be a strong host of mycorrhizal fungi (Soti et al., 2016). Given these benefits, improving the nodulation of sunn hemp by adopting effective inoculation technique would ensure more nitrogen fixation and improve overall soil health.

The objectives of this study were 1) to analyze the effectiveness of different sticking agents in aiding inoculation of *Bradyrhizobium japonicum* L. in sunn hemp seeds to enhance root nodule formation, 2) to observe nodulation pattern over time as affected by the treatment and, 3) to check significant differences between main root and lateral root nodulation pattern due to the treatments. For this purpose, we conducted two field studies, in fall 2019 and summer 2020 to assess the efficiency of different sticking agents peanut oil, jaggery, gum arabic and water (control) on nodulation in sunn hemp in an organically managed field in south Texas.

## 2. MATERIALS AND METHODS

### 2.1 Plant material, adhesive agents, and seed inoculation

Certified organic seeds of sunn hemp ‘Tropic Sun’ were purchased from Johnny’s Seed Co, Winslow ME. The seeds were planted at a rate of 28 kg/ha with each treatment receiving approximately 185g of seed. The sticking agents used were water (1ml/200g seed), peanut oil (1ml/200g seed), gum arabic (40g/100ml), and jaggery(10g/100ml). Seeds in each treatment were weighed into separate polythene bags where they were mixed with their respective adhesive. Powdered peat-based inoculum containing *Bradyrhizobum japonicum* concentration at 2 × 10^8^ CFU/g was used for inoculating the seeds. Approximately 9g of powdered inoculum was weighed and added to each of the polythene bags and shaken vigorously for one minute. After mixing, the contents were spread on a tarp for the coated seeds to air dry for 15 minutes. After the seeds had been dried completely, they were sown using the Jang Seeder (Johnny’s Seed Co, Winslow ME). Seed hoppers were detached and wiped clean before sowing the next treatment.

### 2.2 Site characteristics and experimental design

This study was conducted at the University of Texas Rio Grande Valley (UTRGV) Agroecology Research Garden in Edinburg, TX with a subtropical semi-arid climatic condition. Soil texture in the site was loamy sand with 82% sand, 12% silt, and 6% clay and slightly alkaline pH 7.7. Soil organic matter was 3.8%, total C was 1.2%, and total N was 0.1%. The experiment was a randomized complete block design with four sticking agent treatments (Table 1) having seven replications across 28 individual plots or experimental units. A 223m^2^ block at the Agroecology garden was divided into 28, 4.6m^2^ plots with 0.6m distance between each block. The field was irrigated using sprinklers for one hour/day for the first 35 days for proper establishment of the plant root system at initial stage. No fertilizers were added to enhance the plant growth. The fall trial was conducted from 18^th^ September 2019 to 20^th^ November 2019 and the summer trial was conducted from May 19^th^ to 20^th^ July 2020. The climatic conditions for the experimental site during the two study periods are given in Table 2.

**Table 1:** Concentrations and sources of the treatments used as coating material.

| Treatment type | Coating Material | Concentration | Source |
| --- | --- | --- | --- |
| T1 | Water (control) | 1ml/200g seeds | Tap water |
| T2 | Gum arabic | 40g/100ml | Pure organic ingredients brand |
| T3 | Peanut oil | 1ml/200g seeds | Spectrum Organic |
| T4 | Jaggery | 10g/100ml | Pure Indian foods brand (USDA organic) |

**Table 2:** Meteorological Average of the experimental site during 2019-2020

| Month | Temperature (°C) |  | Rainfall (mm) | Humidity (%) |
| --- | --- | --- | --- | --- |
|  | minimum | maximum |  |  |
| September 2019 | 24.0 | 32.8 | 3.7 | 69 |
| October 2019 | 20.8 | 30.5 | 2.4 | 65 |
| November 2019 | 16.5 | 26.3 | 1.5 | 65 |
| May 2020 | 23.2 | 34.3 | 2.3 | 64 |
| June 2020 | 25.2 | 35.1 | 2 | 64 |
| July 2020 | 25.5 | 35.5 | 2 | 64 |

### 2.3 Data collection

One plant was randomly sampled from each of the plots biweekly and examined for evidence of nodulation and their position on the roots. Since the plants were not densely planted, a circle was marked on surface of soil which approximately included the area of the root underneath. The plant was then gently uprooted with the soil intact to minimize the loss of nodules. Nodules were counted after gently removing the soil from the roots by hand and the position of nodules were also noted. Nodules were investigated on the main root and lateral roots. After removing the nodules separately for main and lateral root, they were washed with tap water and taken to the lab for visual inspection of active nodulation. Nodules were splicing into two halves and checked for reddening, a sign for presence of live nodules containing leghemoglobin and the data for active nodules were recorded.

During sowing and on every nodule data collection day, soil moisture was measured using Digital Soil Moisture Meter with 20 cm probe and soil temperature was measured using Rapitest Digital Soil Thermometer. This process was continued, and the data recorded until sunn hemp started producing flowers. Comparisons across means was done on nodules within weeks and across weeks. Since the total number of nodules produced were different across treatments, percentage of active nodules in each treatment was calculated across weeks.

### 2.4 Data Analysis

For each of the treatments, gum arabic, oil, jaggery and water, a one-way ANOVA was conducted to compare the main effects of each treatment on number of nodules, active or inactive nodules, position of nodules in the root section, percentage of active nodules across weeks and percentage of lateral nodules across weeks. All our tests were two-tailed, performed at a significance level of 0.05 using R version 3.6.2 (R foundation for Statistical computing, Vienna, AT). Normality checks were carried out using Q-Q plot and our assumptions met. For post hoc analysis of means that were significantly different, Tukey’s HSD (Honestly significance test) post hoc analysis with studentized range statistic was done.

Nodulation percentage of each treatment was calculated as the ratio of active nodules by total nodules across each treatment and percentage lateral root nodulation was calculated as the ratio of lateral nodule by total nodule across each treatment. The trapezoid method of AUDPC (Madden et al., 2007) was employed to calculate AUNC (area under nodulation curve) to check the intensity of nodulation across time scale (weeks).

## 3. RESULTS

### Performance of Sticking agents

No differences in the total number of nodules across treatments was observed in fall experiment, whereas significant differences in the total number of nodules across treatments was seen in the summer field experiment (Fig 1). In summer, jaggery treatment had highest total number of nodules average mean of 21.21 followed by oil treatment at average mean 17.21. Gum arabic treatment and water had similar average means of 14.14 and 12.78, respectively. But significant differences in percentage of active nodules fixing nitrogen was not observed in summer with active percentage of nodules for jaggery, oil, gum arabic and water treatment as 93%, 90%, 84%, and 84% though differences in total nodules was seen across treatments (Fig 2). However, in fall, differences were seen in percentage of active nodules among the different treatments (P<0.001) over the study period (Fig 2). Water treatment performed least with low nodulation and low active percentage of only 40% (P<0.001) followed by gum arabic which had better nodulation but only 72% of nodules were active. Jaggery and oil treatment had similar higher active nodules percentage of 87% and 90%, respectively (Fig 4a). Jaggery exhibited higher lateral root nodulation and gum arabic exhibited higher main root nodulation in fall. Overall main root and lateral root nodulation was consistently higher in jaggery followed by oil in summer (Fig 4b).

**Figure 1:**
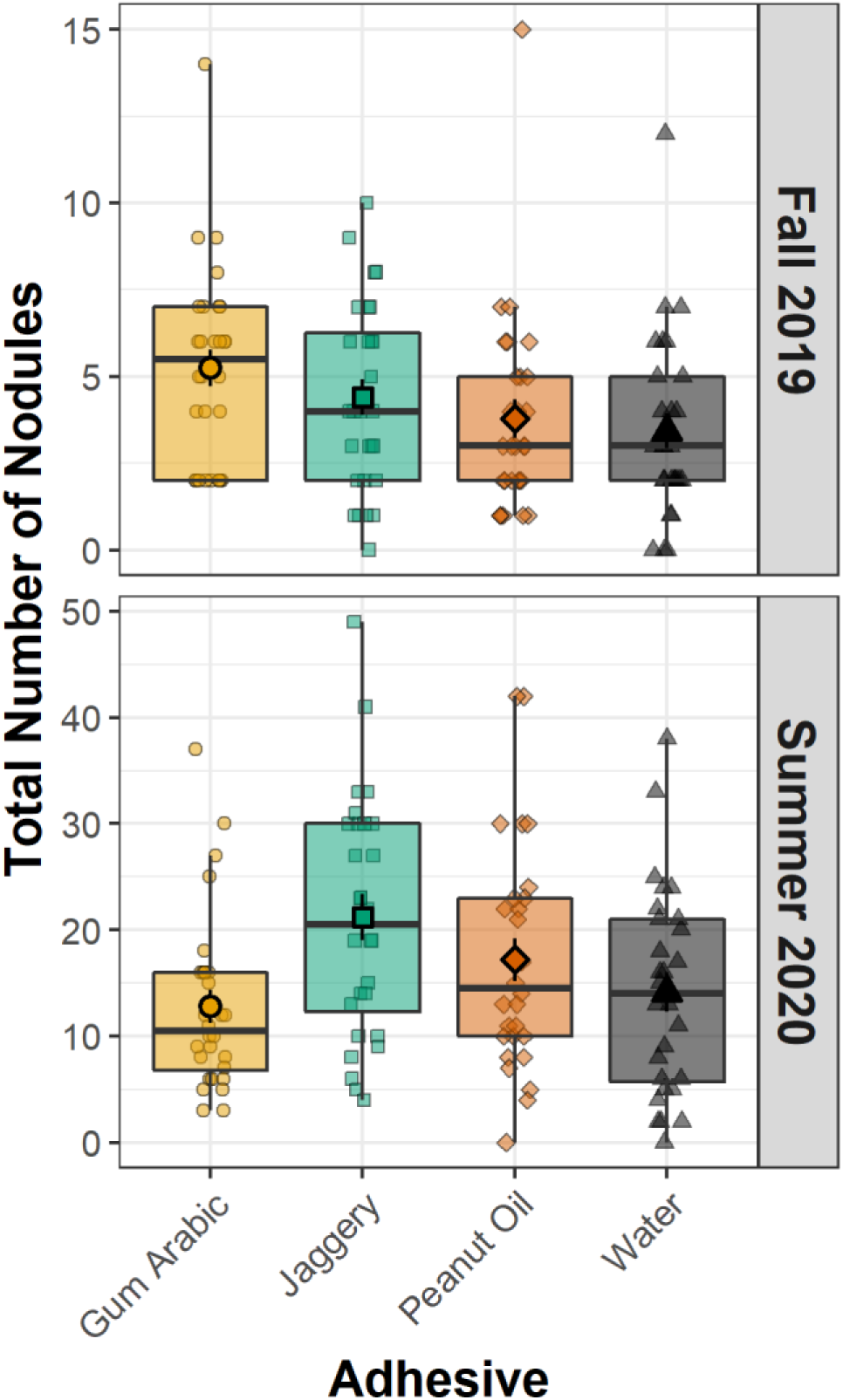
Total number nodules across the different treatments in fall 2019 and summer 2020.

**Figure 2:**
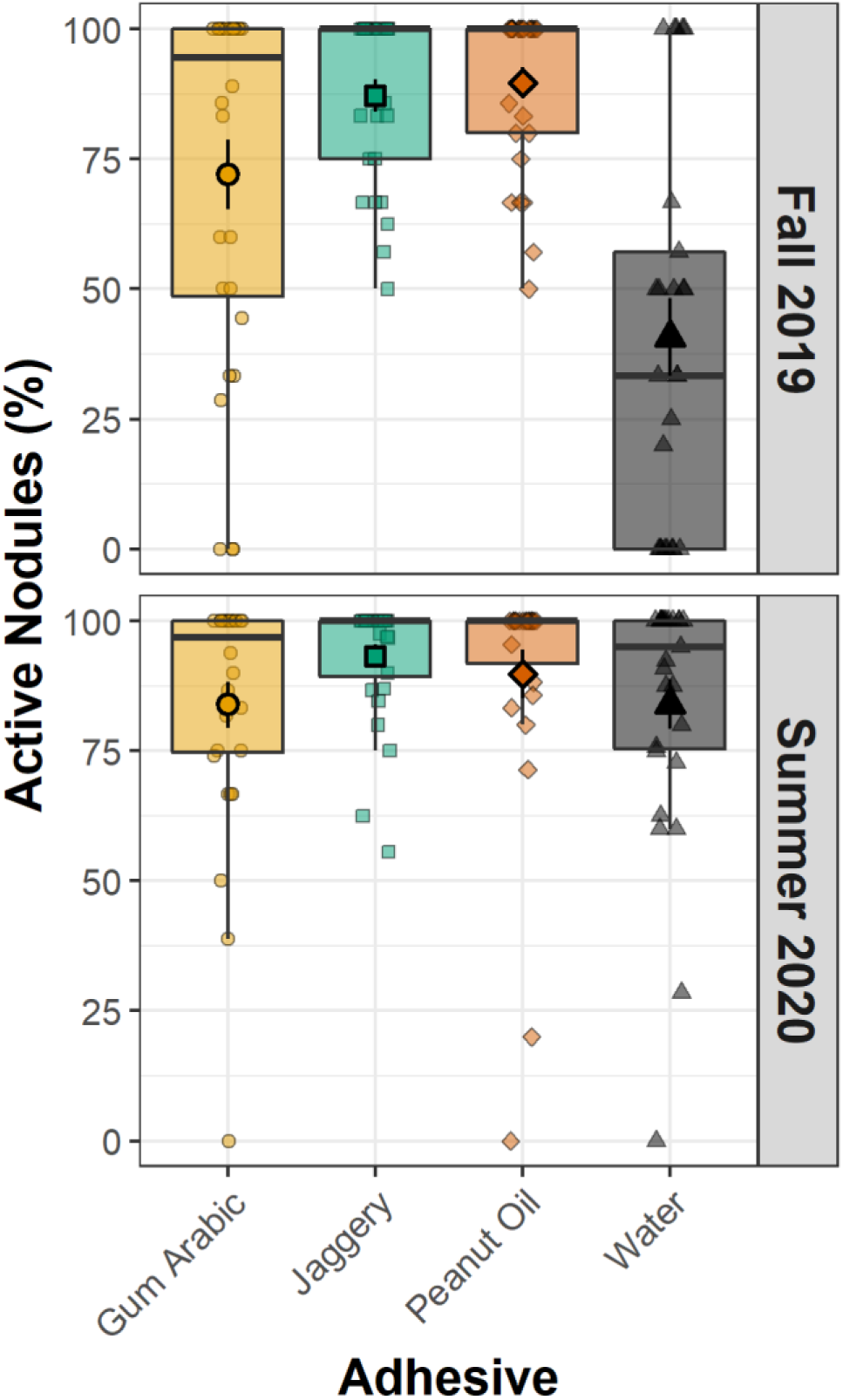
The boxplots show the total mean active nodulation (N=28) throughout the experiment across the treatments in the fall 2019 and summer 2020 field study.

### Nodulation over time

Nodules were collected after 2, 4, 6, and 8 weeks after sowing and data recorded for total nodule number, activity of nodule, and position of nodule. Percentage of active nodules significantly changed over time in the fall experiment (Fig 3). At week 2 jaggery treatment had the highest percentage of active nodules at 100% followed by oil at 90.9%, while the water treatment, control, had the lowest percentage of active nodules at 70%. At week 4, there was a decline in the number of active nodules, with the water treatment having the largest decline of 39%. The peanut oil treatment had consistently high percentage of active nodules with highest nodulation later in week 6 at 93.75% and week 8 at 87.5% subsequently. Gum arabic treatment had 85%, 75%, and 91% nodulation at 2, 4, and 6 weeks respectively and showed a sharp decline in active nodule at week 8 which lowered the total active nodule percentage to 33.33%. Water treatment showed a consistent decline of total nodules and percent of active nodules over the study period.

**Figure 3:**
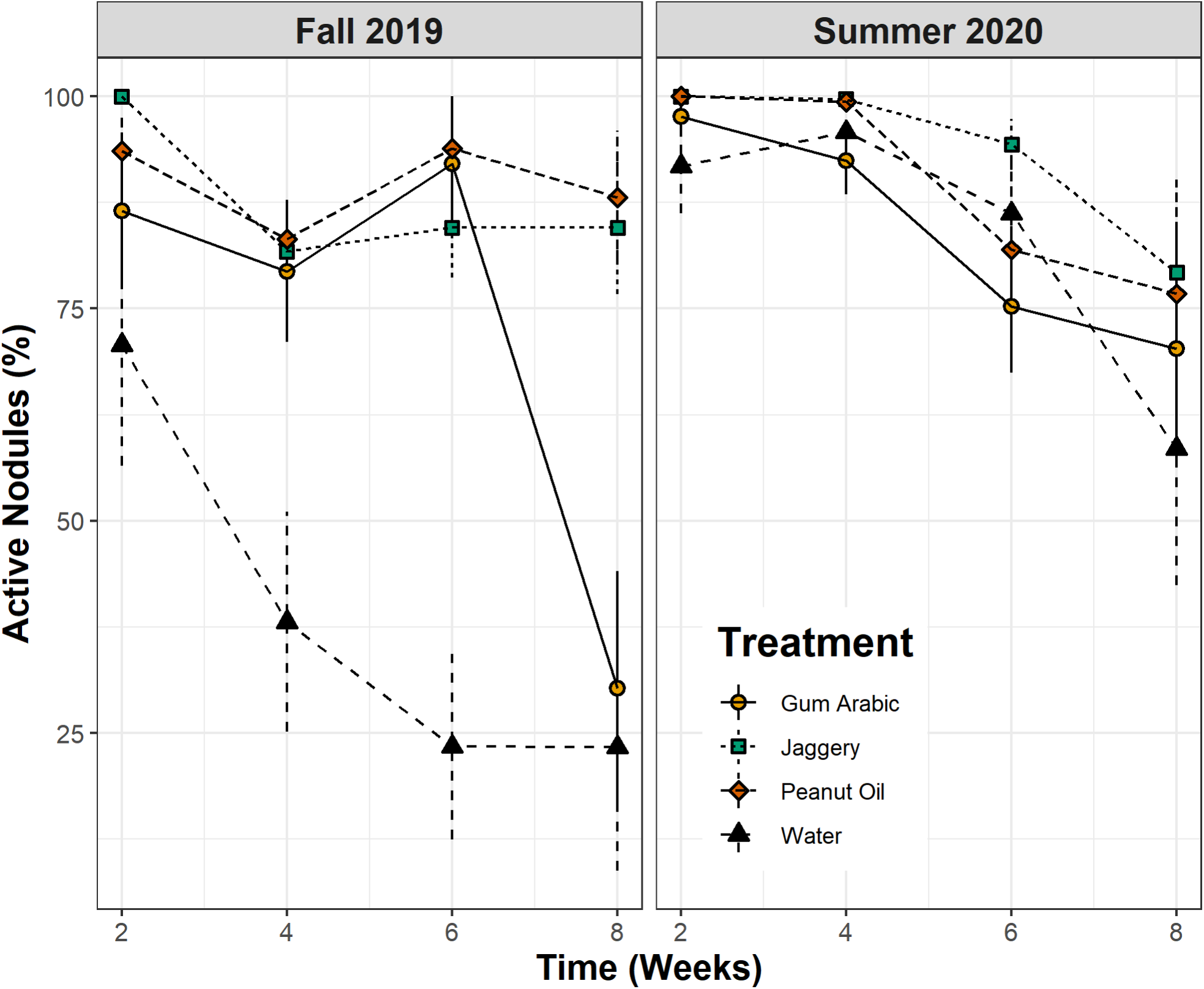
The line graph shows the pattern of percentage of active nodules for weeks 2, 4, 6 and 8 in the fall 2019 and summer 2020 field studies.

In summer the nodulation declined over time for all treatments (Fig 3). But the overall nodule number was at least three times larger than the fall study for all treatments. Oil and jaggery had ∼100% nodulation in week 2 and 4 with higher active nodulation percentage as well. Gum arabic and water treatment had no significant differences in nodulation across weeks (p<0.05) and had active nodulation of 83% for both. Week 2 and week 4 exhibited the best performance of jaggery in both lateral and main root nodulation and higher active nodulation (p<0.001) followed closely by oil treatment. Area under nodulation curve (AUNC) for percentage of active nodules value for oil, jaggery, gum arabic and water treatments were 536, 491, 460, and 240 for fall with significant lower value for water and peanut oil with the highest for fall. Jaggery showed highest AUNC value of 567 for percentage active nodules, and peanut oil, water, and gum arabic had AUNC value of 537, 513 and 503, with gum arabic showing least performance in summer.

### Main versus lateral root nodulation

Nodulation pattern in gum arabic had a highest mean for main root nodulation and jaggery had a significant higher lateral root nodulation mean over the weeks in fall (Fig 4a). Water treatment had the least lateral root nodulation over the weeks. However, in summer, jaggery treatment had overall higher main root and lateral root nodulation mean closely followed by peanut oil treatment (Fig 4b). There was no significant difference in the soil moisture 20.9% and soil temperature 28.2 °C across treatments in both the experimental study seasons.

**Fig 4a:**
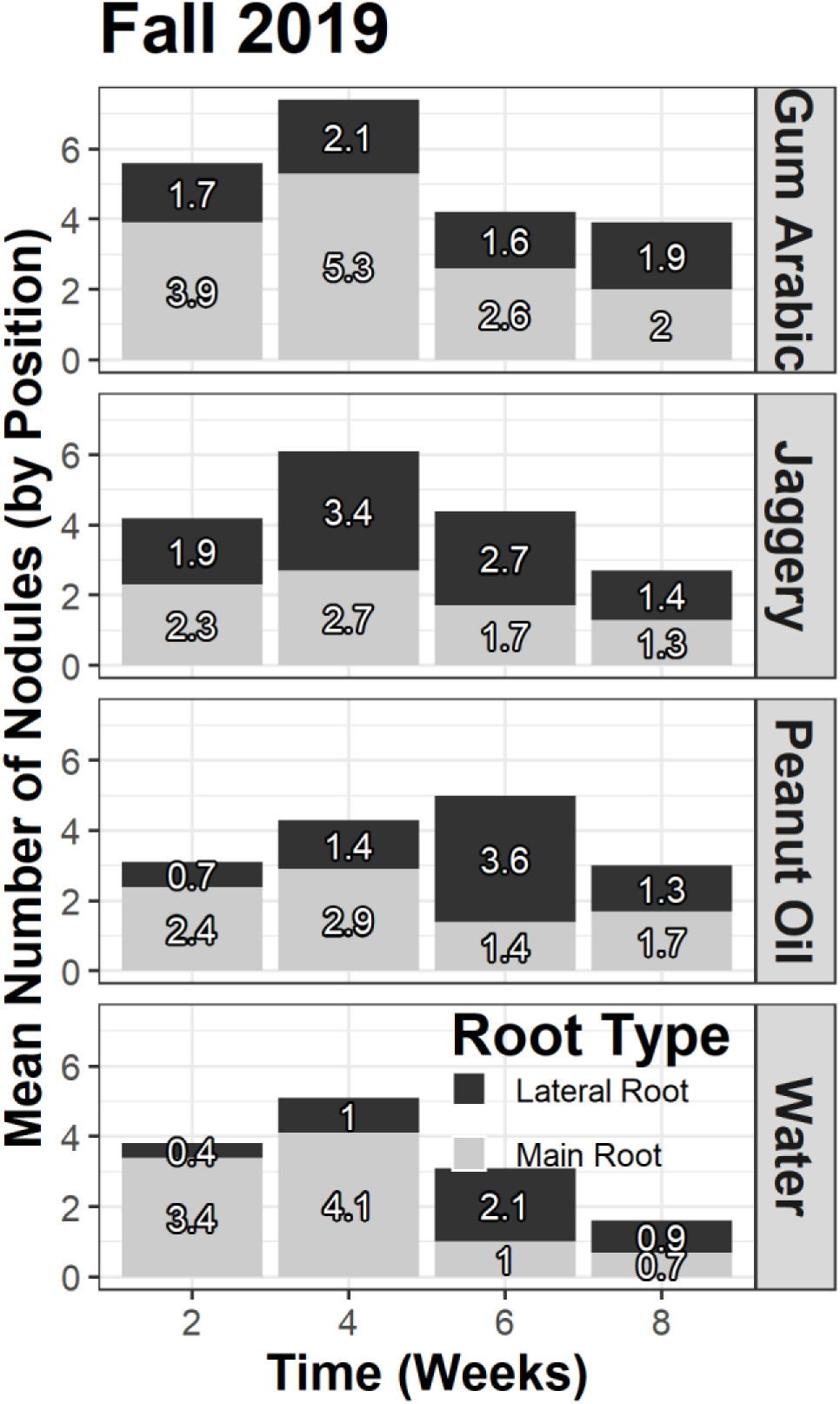
Facet grid plot illustrating the differences in the position of nodules in roots for each treatment (n =28) across the eight weeks during the fall 2019 field study.

**Fig 4b:**
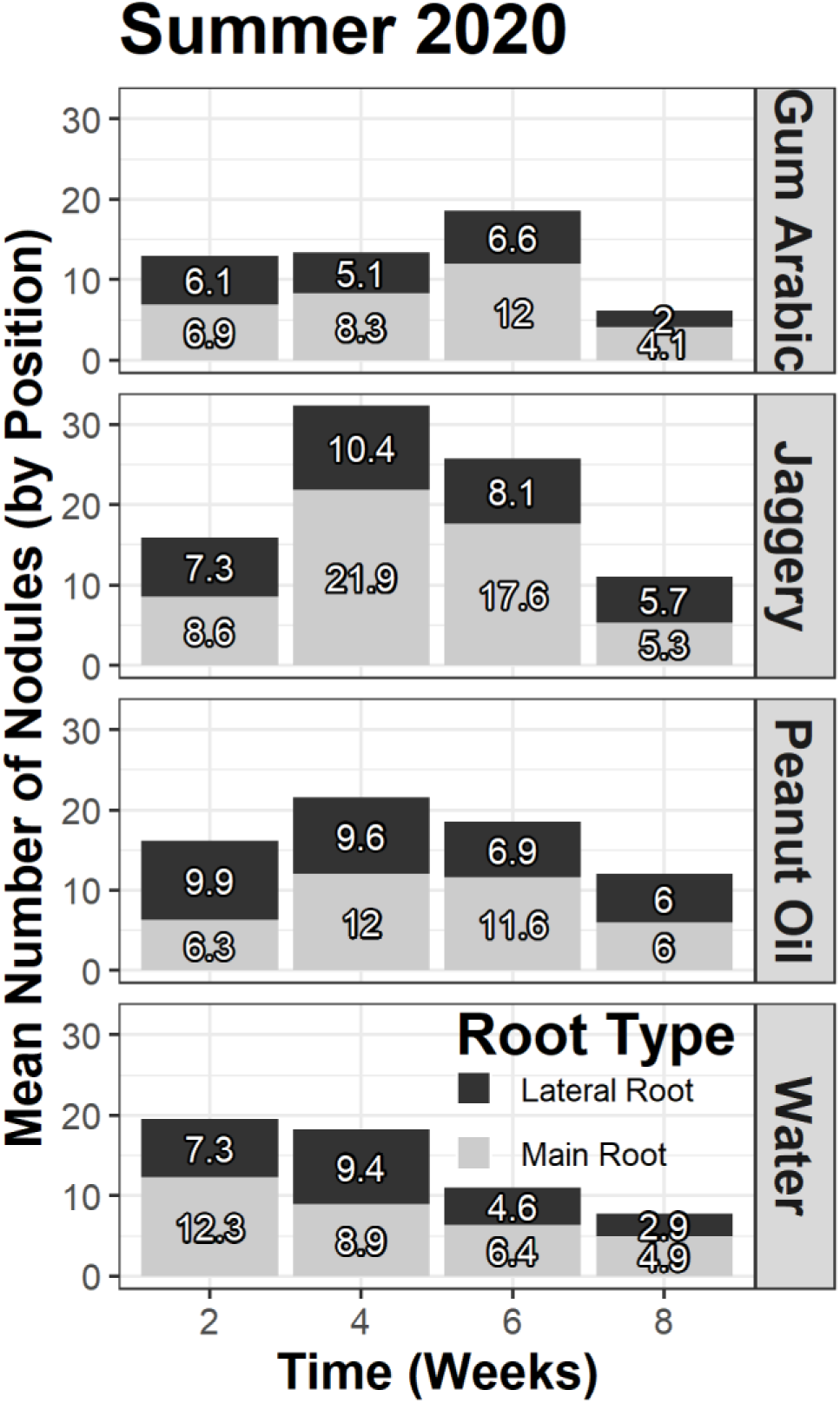
Facet grid plot illustrating the differences in the position of nodules in roots for each treatment (n =28) across the eight weeks during the summer 2020 field study.

## 4. DISCUSSION AND CONCLUSIONS

The purpose of using sticking agents in seed inoculation is to ensure the close contact with seeds and the survival of the inoculants. In this study we analyzed nodulation in a popular leguminous summer cover crop, sunn hemp, as affected by use of adhesives for *Bradyrhizobium* inoculation at sowing and other characteristics on nodule formation over two seasons, fall 2019 and summer 2020. We explored the potential effects of the four commonly used sticking agents: water (control), gum arabic, peanut oil, and jaggery. While there were slight differences between the two growing seasons, overall gum arabic, peanut oil, and jaggery treatment did significantly better than water treatment.

Gum arabic held higher total number of nodules in the main root during the fall study. The number of nodules declined as weeks progressed in gum arabic treatment and ratio of active nodules was less compared to the nodules in both seasons. Gum arabic, a gum derived from exudates of *Acacia senegal* is an effective adhesive able to bind the inoculant well and produce maximum nodules (Elegba and Rennie, 1984; Swarnalakshmi et al., 2011). Due to its polymeric nature, gum arabic could maintain the viability of rhizobacteria on legume seed and protect them from desiccation. Previous *in vitro* experiment also shows, gum arabic was more tenacious and provided good adhesion holding almost equal number of rhizobacteria compared to jaggery (Date, 1970; Swarnalakshmi et al., 2011).

However, in our study jaggery performed better than gum arabic treatment or water treatment getting highest main root nodulation during fall with 87.23% active nodulation followed by highest main and lateral root nodulation in summer trial as well. Jaggery is an unrefined sugar produced by evaporating water from sugarcane juice and widely used for seed inoculation by the Indian farmers (Sayyed et al., 2012; Swarnalakshmi et al., 2011). Previous study revealed local *Rhizobium* strain UR5 inoculation on urd bean using jaggery as carrier significantly resulted in higher number of nodules and the trend was similar in consecutive three years from 2008-2011 when compared with other carriers like lime pelleting, lignite, sawdust, and sterilized cowdung powder (Rajbhoj and Baig, 2011). However, another research shows, jaggery initially resulted in good coating but later became brittle pellet after evaporation and absorption into the seed. Also, survival of rhizobia with carboxy methyl cellulose and gum arabic was found better in comparison to jaggery carrier (Swarnalakshmi et al., 2011). For, peanut oil treatment, consistent higher active nodules were observed in both seasons even though total number of nodules were similar to water treatment. Water treatment performed least resulting in lesser nodulation with lesser active nodules which is in accordance with other similar results (Elegba and Rennie, 1984; Hoben et al., 1991). Apart from differences in total nodules and total active nodules, some treatments also have exhibited inclination towards main root or lateral root nodulation in the root system.

Gum arabic held higher total number of nodules in the main root around the crown area in the fall study. Previous research shows, crown area nodulation contained superior *Bradyrhizobium* from the inoculum in soyabeans since this bacterium has limited mobility and restricted to root crown area where it was initially placed during planting (McDermott and Graham, 1989; Wadisirisuk et al., 1989). Even the Food and Agriculture Organization of the UN widely held opinion was that that nodules clustered on main root are more effective than nodules on lateral roots (Alaa El Din et al., 1984). Since gum arabic is a strong adhesive and may have populated *Bradyrhizobium* at main root, there can be a possibility of maximum nodulation with maximum nitrogen fixation through use of gum arabic as sticking agent.

Some other research on position of nodulation has conflicting views. Nodulation on *Glycine max* (L.) Merrill records nodules formed on lateral roots and at a greater soil depth contributed more to nitrogen fixation during the growing season than those formed on tap root or 1-5cm soil depth (Hardarson et al., 1989). In our experiment, the jaggery treatment seemed to be a better sticking agent in the later stage of nodulation where it develops lateral roots in both seasons compared to rest. Lateral root nodules are important in providing fixed N during latter stages (McDermott and Graham, 1989). Weaver and Frederick (1974) found that inoculation increased nodulation on lateral roots in soils containing very less rhizobia in soil. Since our experimental field was not used for any agricultural purpose previously, they may not have or have very less indigenous *Bradyrhizobium*. Good sticking agents help successful nodulation and choice of sticking agent may even help the pattern in which nodulation in root system takes place according to our experiment. Choosing of sticking however needs to be employed by considering the ease of its applicability, availability, costs, and other aspects.

Considering application and availability of sticking agent, water was the easiest to apply and gave an even layer of inoculum around seed. But, after drying muchof the inoculum started falling off the seed coat. Even some amount of inoculum remained in the seed hopper after seeding. Regardless, water is readily available as a sticking agent and does not incur significant extra costs to the farmer. In contrast, gum arabic is expensive and not as readily available, but can be an extremely good adhesive under certain circumstances. It can however be strong enough to even bind seeds together forming a clump causing difficulty in using the seed hopper adjusted to the rate of sunn hemp as per the experiment. Gum arabic 40% solution is the recommended adhesive mixture for peat inoculum (Date, 1970). A thinner gum arabic solution can be something to look during future adhesive use. Peanut oil facilitated easy application and did not clog the pores on the seed hoppers. It is not as good an adhesive as gum arabic and managed to stick only 70% of the inoculum. Peanut oil is readily available cooking oil in many households and not very expensive as well as does not attract ants and other insects, unlike some of the sugary adhesives like jaggery and molasses. The jaggery treatment did not attract ants, despite being primarily composed of sugar. Though Jaggery was sticky to mix, once it dried with the inoculum, it formed stable pellets that could be easily sown using the seed hoppers. Though jaggery is a household item in most Asian countries like India (Swarnalakshmi et al., 2011), it may not be available in other countries.

The area under the nodulation curve (AUNC) is the estimated quantitative summary of nodule intensity over time, for comparison across weeks. For estimating the AUNC, the trapezoidal method was used which discretized the time variable (weeks) and calculated the average nodulation intensity between each pair of adjacent time points. The nodulation intensity for percentage of active nodules in fall study for four treatments was significantly lower for water treatment at 240, suggesting the lowest intensity of active nodulation among others. Oil treatment at 536 suggested highest active nodules, closely followed by jaggery treatment, and gum arabic treatment at 491 and 460, respectively. In summer, jaggery treatment had the highest active nodulation intensity at 567 closely followed by oil at 537 and gum arabic had the lowest AUNC value of 503 suggesting poor nodulation intensity. These values suggest oil and jaggery may be better suitable for as treatments for increased intensity of active nodules. Though AUNC value is usually not applied to estimate nodulation, it may be a useful tool to quantify the progression of nodules in future studies.

Apart from differences in nodulation among treatments within the season, it was seen that nodulation in summer 2020 was approximately three times higherthan in fall 2019 for all the treatments. Sunn hemp is said to grow well in warmer weather conditions, putting better biomass and delayed flowering (USDA-NRCS 2005). Although differences due to soil temperature in nodulation across treatments were not seen, the summer nodulation increase can be explained by the fact that the population of *Bradyrhizobium* in the soil may have improved, due to fall planting with inoculation followed by successful sunn hemp nodulation. Also, understanding phytomicrobiota around the root nodule region is important, as the interaction among individual in a microbiome is complex, sometime even affecting the behavior and fitness of host plant and often aiding or repelling nodule formation (Martínez-Hidalgo and Hirsch, 2017). The phytomicrobiota may contain arbuscular mycorrhizal fungi which is seen to help nodulation and the symbiotic association to form nodules, gram-negative bacteria like *Pseudomonas, Kliebsiella, Ochrobactrum* and gram-positive bacteria like *Bacillus, Paenibacillus, Lysinobacillus*, and Actinomycetes like *Micrpminospora, Streptomyces* and Nitrogen-fixing *Frankia* (Martínez-Hidalgo and Hirsch, 2017). Significant increase in nodule biomass was observed when *Pseudomonas* strains were co-inoculated with the *Mesorhizobium* sp. *Cicer* strain Ca181 and *Pseudomonas* (Sindhu and Dadarwal, 2001). The dynamic of the soil and the microbiota could have been altered by cultivation of sunn hemp and *Bradyrhizobium* inoculated. Indigenous soil strains population alteration may also be responsible for the nodulation sometimes.

It is often noted that inoculant *Bradyrhizobia* show competition with indigenous soil strains. However, they are restricted at the point of placement in the soil, with limited mobility they are unable to spread throughout the developing root system (Vlassak et al., 1997). This could often result in low nodule occupancy by the inoculant strain in the tap and lateral roots. Further research on the reasons that may have resulted on enhancing lateral or crown root nodulation may be a potential avenue. Checking nitrogen in the nodules based on position and timing of nodulation is also encouraged. More research is needed on the differences between lateral and main root nodulation and how rhizobia populations associated with different root types may impact overall nitrogen fixation. The benefits and impact to soil fertility can be better understood properly only through multi-years of observation. It would be interesting to plant the plot with sunn hemp and observe over 2-3 years using adhesive based inoculation technique. Also, instead of using peat-based inoculum if researchers used pure *Bradyrhizobium* taken directly from nodules rather than peat and mixed with adhesives could also be an option. Also, other adhesives which have been used before can be explored like carboxymethyl cellulose, wallpaper glue, corn syrup, evaporated milk, honey, and powdered milk (Deaker et al., 2004; Elegba and Rennie, 1984; Rajbhoj and Baig, 2011), which are all readily available in many subtropical regions.

Most research considering sticking agents and rhizobial inoculation are lab experiments usually in pots and closed chambers. Though they may give statistically significant results, this field trial would give results which can be much similar analysis of actual field settings and can be more suggestive as an option. Considering the recent popularity of sunn hemp as a nitrogen fixing cover crop in subtropical areas, research on inoculation method of seed using suitable adhesive to ensure live rhizobial adherence till successful nitrogen fixation is useful. Also, there is not much information on lateral and main root nodulation for sunn hemp, or nodulation, and activity of nodules. This paper would work as a guide to field inoculation of sunn hemp plantation. Since the area under experimentation was previously never used for agriculture, they may be mostly devoid or have fewer indigenous *Bradyrhizobium* compared with areas that had previously been cultivated, the nodules are a good representative of the inoculum *Bradyrhizobium* added. Since, this research also shows nodulation over time compared across treatments and would give a base to future research aimed at seeing nitrogen fixation at different stages of nodule development as well as based on the position of nodule.

## 5. ACKNOWLEDGEMENTS

This work is done as part of the Subtropical Soil Health Initiative is supported by the Conservation Innovation Grants program of the USDA NRCS, under grant #69-3A75-17-281. Student support came from UTRGV Presidential Graduate Research Assistantship.

